# Encoder-based Curvature-Aware Regularization for estimating asymmetric fiber orientation distribution functions in diffusion MRI

**DOI:** 10.64898/2026.03.31.715534

**Authors:** Mojtaba Taherkhani, Marco Pizzolato, Morten Mørup, Tim B. Dyrby

**Affiliations:** Department of Information Engineering, University of Pisa, Pisa, Italy; Department of Applied Mathematics and Computer Science, Technical University of Denmark, Kgs. Lyngby, Denmark; Danish Research Centre for Magnetic Resonance, Copenhagen University Hospital Amager and Hvidovre, Copenhagen, Denmark

**Keywords:** Diffusion MRI (dMRI), Asymmetric fiber orientation distributions (A-FODs), self-supervised Transformer, Fiber Continuity

## Abstract

Diffusion-weighted magnetic resonance imaging (dMRI) is used to study white matter microstructure and to delineate pathways by estimating fiber orientation distributions (FODs). Symmetric FODs represent the conventional model assuming antipodal symmetry in water diffusion. However, in complex regions with bending, branching or fanning fibers, this assumption is not guaranteed. To better capture such underlying fibers geometries, asymmetric FODs (A-FODs), derived from neighboring FODs, have been introduced. Here, we propose an Encoder-based Curvature-Aware Regularization (EnCAR) method for estimating A-FODs. Incorporating curvature features into the regularization weight applied to neighboring voxels improves reconstruction of A-FODs. A self-supervised Transformer network, combined with a Spherical Harmonics Semantic Encoder, learns region-specific regularization parameters from this local neighborhood to capture the diversity of fiber geometries across the brain. The EnCAR method was verified on the DiSCo challenge phantom, and applied to in vivo multi-shell Human data. The model estimated sharp, high-angular-resolution A-FODs that were well aligned with local fiber pathway. Compared with established FOD and A-FOD methods, it performed on par in regions dominated by symmetric FODs and outperformed them in complex asymmetric regions. Quantitative evaluation using the Asymmetry Index (ASI) and Model Discrepancy Index (MDI) confirmed improved consistency with the underlying diffusion signals. By ensuring smooth directional transitions, this work enhances the visibility of continuous fiber segments.

## 1. Introduction

Diffusion-weighted magnetic resonance imaging (dMRI) is a non-invasive quantitative modality that enables investigation of the brain’s anatomical organization in vivo at the macroscopic scale [1–3]. This technique allows extracting voxel-wise fiber orientation distributions (FODs) by characterizing the anisotropy of water diffusion, particularly in white matter containing densely packed axons [4, 5]. Since dMRI cannot distinguish between opposing diffusion directions, conventional FODs are represented as antipodally symmetric.

While symmetric FODs are widely used in white and gray matter, they are limited to capturing symmetric single or multiple fiber populations [4, 6]. Due to finite imaging resolution, sub-voxel anatomical information of pathway geometry is lost and enforced to a symmetric FOD [7]. In regions with bending, branching, or fanning fibers, the true FOD may be asymmetric, forming T- or Y-shaped patterns, distinct from symmetric crossing fibers at the sub-voxel level.

Asymmetric FODs (A-FODs) have been proposed to recover such sub-voxel information using neighborhood FODs [4, 8–11]. A-FODs improve alignment with complex local pathway organizations, such as crossing and bending fibers, and improve fiber tractography compared to approaches based on symmetric FODs [2, 8, 11]. One class of methods, commonly referred to as filtering-based approaches, generates A-FODs after conventional FOD reconstruction by applying regularization across neighboring voxels. Regularization weights are defined using inter-voxel distance, directional alignment, orientation similarity, and intensity similarity, modeled using distributions such as Gaussian and von Mises–Fisher and combined into a unified weight [4, 8, 10, 12].

A central assumption in these methods is the principle of fiber continuity. This principle posits that neural fibers typically propagate along nearly straight, continuous trajectories between voxels. Early approaches applied this principle in different ways. For example, [12] iteratively refined asymmetric spherical functions using displacement probability, inter-voxel distance, directional alignment, and orientation similarity. CB-REG [10] emphasized neighborhood structure by dividing voxels into forward and backward groups and applying separate Gaussian kernels. Other methods [4, 8, 13] incorporated filtering, cone models, or constrained spherical deconvolution to enhance asymmetry reconstruction and mitigate biases such as the gyral bias[5, 9].

Despite these advances, current methods share two main limitations. First, the assumption that fibers propagate as straight lines between the central voxel and all neighboring voxels may not hold as the spatial distance increases, reducing reconstruction accuracy [7]; consequently, it is necessary to incorporate the curvature characteristic into the regularization. Second, the use of global regularization parameters, where a fixed set of parameters is applied to all voxels, fails to capture regional variations in fiber geometry. A strategy that adapts regularization locally is therefore needed to better reflect the underlying geometry.

The aim of this work is to propose the Encoder-based Curvature-Aware Regularization (EnCAR) method, which functions as an adaptive continuity model, overcoming the strict straight-line constraint to generate accurate A-FODs. A self-supervised Transformer network learns region-specific regularization parameters to capture local structural variations. To enhance its ability to interpret complex geometric relationships, we introduce the Spherical Harmonics Semantic Encoder (SH Semantic Encoder) as an input embedding layer, which resolves ambiguities in raw SH coefficients and reduces model complexity. The architecture projects semantic relationships between neighboring voxels into the parameter space of the regularization weights, enabling the reconstruction of A-FODs consistent with local fiber geometry.

The network is trained in a self-supervised manner by minimizing the mean squared error (MSE) between input FODs and the symmetric component of the estimated A-FODs. In this study, we hypothesize that EnCAR, through its enhanced curvature awareness, can more accurately resolve complex A-FODs well aligned with local fiber pathway. To investigate this, we compare our framework against existing FOD and A-FOD methods [8, 12], validating its performance on the DiSCo challenge phantom [14] and a Multi-shell human dataset [15].

## 2. Methods

In the general formulation for filtering-based methods[8], A-FODs are modeled as a weighted combination of symmetric FODs within a local neighborhood:

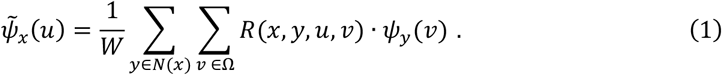

Here, 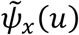 denotes the A-FOD at voxel position *x* along direction *u*, and *N*(*x*) represents the local neighborhood of voxels surrounding the central voxel *x*. The set *Ω* corresponds to the unit sphere directions onto which the neighboring FOD is projected. W is the normalization factor that ensures correct scaling of the resulting A-FOD (center voxel), and *R*(*x, y, u, v*) represents the regularization weight applied to the neighboring symmetric FOD *ψ*_*y*_(*v*) at voxel *y* in the direction of *v*. Furthermore, the displacement vector from the center voxel *x* to the neighboring voxel *y* is decomposed into its magnitude *I*_*xy*_ and unit direction vector *D*_*xy*_. Our work extends this general formulation by introducing a novel, curvature-based regularization weight, *R*_*θ*_(*x, y, u, v*), whose parameters are learned in a region-specific manner by a Transformer model.

### 2.1. Curvature-Aware Regularization

We introduce a curvature-aware regularization designed to reflect fiber curvature from neighboring voxel toward the center voxel when estimating the A-FOD. The regularization weight is constructed by employing three key factors: inter-voxel distance, directional alignment, and orientational similarity.

The necessity for incorporating curvature properties is visually illustrated in Fig. 1a. When the spatial distance between the central voxel and its neighboring voxels becomes relatively large, the straight-path assumption does not hold in all situations [7], as the distance between voxel centers may exceed the radius of curvature of the fiber.

**Figure 1.**
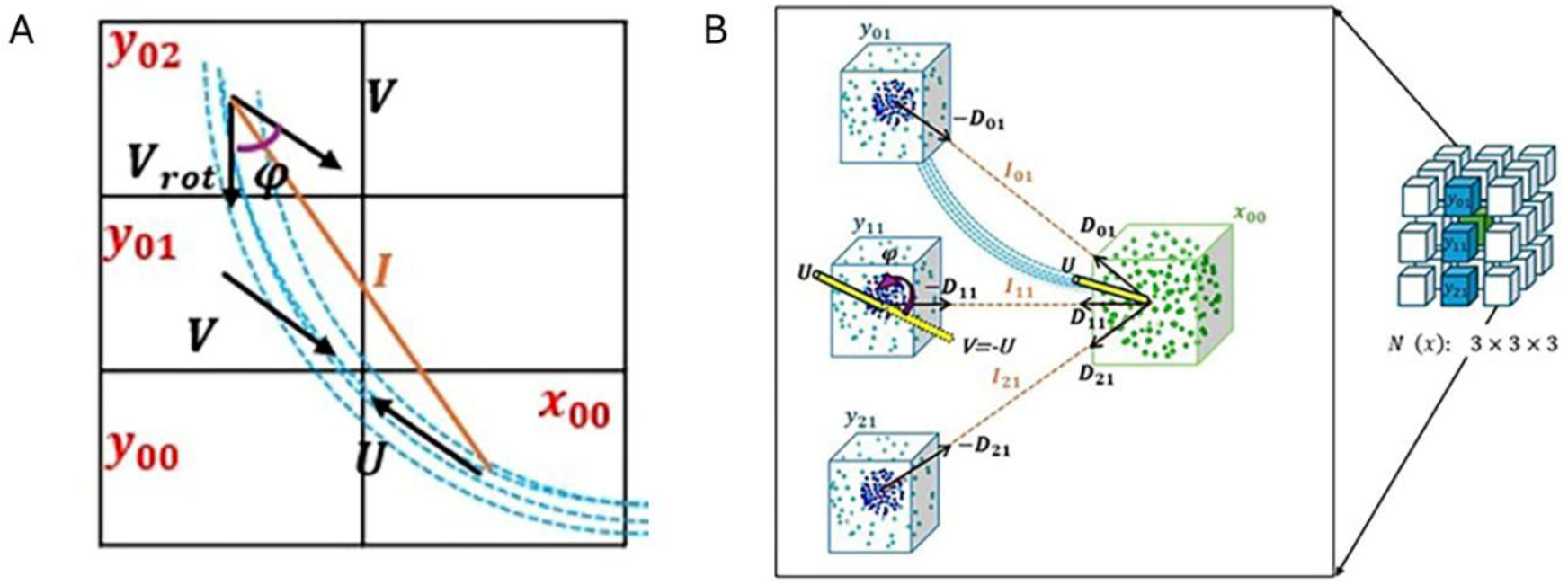
Visual interpretation of the curvature-aware regularization prarameters, central voxel x and its neighbors y. (A) A 2D schematic of a curved fiber pathway illustrating the neighbor’s original direction V and rotated-adjusted direction to account for the fiber’s curvature. (B) A view of the local neighborhood illustrating key parameters bettween the center voxel and selected voxels.

To embed curvature information into the regularization weight, the direction *υ* at neighboring voxels used in orientational similarity is rotated along the vector *D*_*xy*_. The rotation is modulated by the inter-voxel distance *I*_*xy*_ via a weighted polynomial arc function, defined as

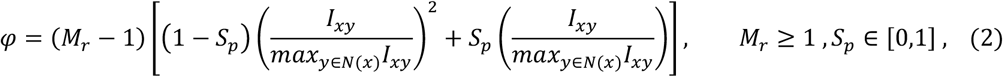

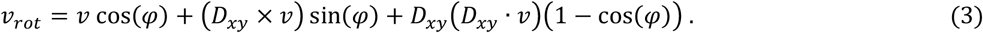

Here, *v*_*rot*_ is the rotated direction, calculated using Rodrigues’ rotation formula [16], where *φ* represents the rotation angle. The parameter *S*_*p*_ determines the shape of the arc, while *M*_*r*_ defines the maximum allowable rotation. Specifically, *M*_*r*_ = 1 corresponds to no rotation. Regarding the path, *S*_*p*_ = 1 produces a trajectory with constant curvature, whereas *S*_*p*_ = 0, yields a quadratic curve path. The arc formulation ensures that rotation initiates at zero at the center and smoothly accelerates to a maximum at the boundaries of *N*(*x*), thereby enabling the framework to accurately capture curved fiber trajectories. The curvature-aware regularization is defined in Eq. (4):

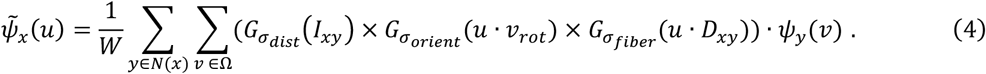

Here, *G*_*σ*_ is a Gaussian distribution with unit mean (*μ* = 1) and standard deviation *σ*. The term 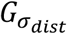 assigns weights to neighboring voxels based on their Euclidean distance from voxel *x*. The function 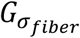 accounts for the alignment between the currently processed direction *u* and direction *D*_*xy*_. Similarly, the 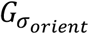 models the alignment between the current direction *u* and the curvature-adjusted direction *v*_*rot*_, derived from Eq. (3). A visual interpretation of the parameters in Eq. (4) is presented in Fig. 1.

### 2.2. self-supervised transformer architecture

The Transformer Encoder [17] is employed to encode the semantic relationships among neighboring voxel set and project them into the regularization parameter space defined in Eq. (4). The Transformer Encoder is selected due to its capacity to model long-range dependencies and capture global contextual patterns across input tokens[18]. Fig. 2 provides an overview of the proposed architecture.

**Figure 2.**
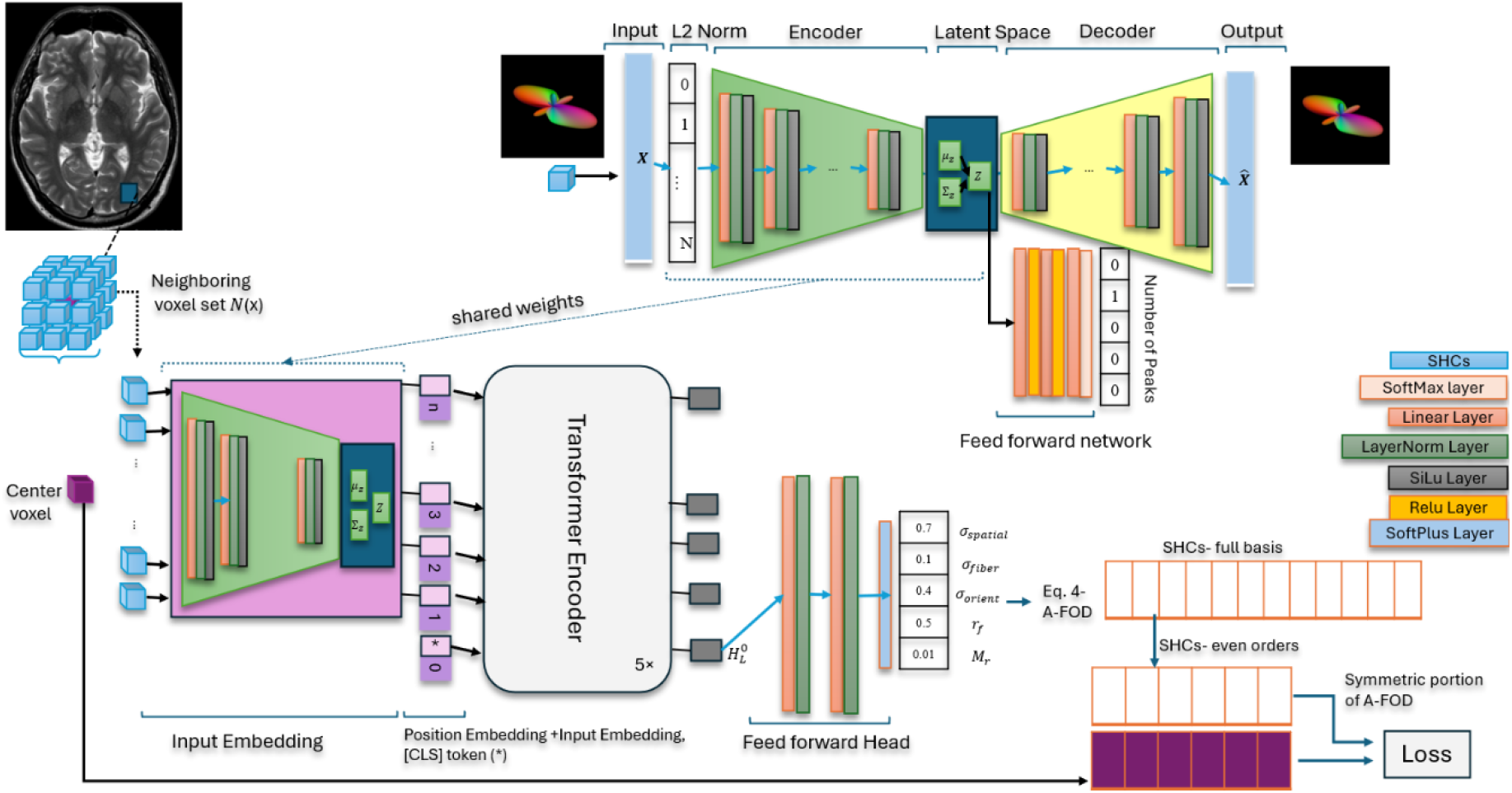
An overview of the self-supervised Encoder-based Curvature-Aware Regularization (*EnCAR*). Symmetric FODs are projected into a semantic latent space by a VAE-based encoder. This sequence of embeddings is processed by a Transformer Encoder and then mapped by a feedforward head to the five parameters of the curvature-aware regularization. These parameters are used to reconstruct the central voxel’s A-FOD. The symmetric component of A-FOD and the original input FOD is used to train the model.

The voxel set *N*(*x*) serves as the input sequence of the network. Within each voxel, the symmetric FOD is represented by its even-order spherical harmonics coefficients (SHCs) [19, 20]. Thus, the input sequence is denoted as 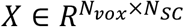, where *N*_*vox*_ is number of voxels in the local neighborhood (sequence length) and *N*_*SC*_ is the number of SHCs per voxel.

Input Embedding (Fig. 2): The standard Transformer expect a 1D sequence of fixed-dimensional token embeddings. However, Different SHC sets can yield similar FOD shapes, introducing representational ambiguity [21]. To mitigate this and enhance semantically meaningful local feature extraction [18], Spherical Harmonics Semantic Encoder (SH Semantic Encoder) is proposed to project the raw SHCs of each voxel into a compact *D*-dimensional latent semantic space, yielding a sequence of semantically rich token embeddings.

Transformer Encoder (Fig. 2): Inspired by architectures such as BERT [22], ViT[23], BLIP [24], and CLIP [25], we prepend a learnable token [CLS], denoted as 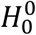, to the input embeddings. Sinusoidal positional embeddings [17] are concatenated to the input embeddings to preserve spatial ordering within the neighborhood. The resulting sequence is then passed through the Transformer Encoder [17] comprising *L* layers, each alternating Multi-Head Self-Attention (MSA) and MLP blocks. LayerNorm is applied before every block, and residual connections after each block [25, 26].

Feedforward Head (Fig. 2): The final representation of the [CLS] token after the last Transformer layer, denoted 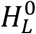, serves as a global aggregated semantic representation of the neighboring voxels. To map the learned feature vector 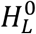 into the regularization parameter space, a feedforward neural network is appended. The architecture consists of a stack of two linear layers interleaved with Layer Normalization (LayerNorm), followed by a final linear mapping with Softplus activation. We refer to this output as the parameter space embedding (*σ*_*dist*_, *σ*_*fiber*_, *σ*_*orient*_, *S*_*p*_ and *M*_*r*_).

#### 2.2.1. Spherical Harmonics Semantic Encoder

To generate the input embedding described above, we employ the Variational Autoencoder (VAE) architecture shown in Fig.2. Using VAE allow us to learn compressed semantic representations from high dimensional data [26]. The encoder part of the pre-trained VAE generates the input embeddings. This VAE is trained to reconstruct a semantic latent space wherein similar FODs (in terms of peak count, profile variance, and peak directions) are mapped to nearby vectors, and dissimilar ones are mapped farther apart.

#### 2.2.1. Spherical Harmonics Semantic Encoder

To generate the input embedding described above, we employ the Variational Autoencoder (VAE) architecture shown in Fig.2. Using VAE allow us to learn compressed semantic representations from high dimensional data [26]. The encoder part of the pre-trained VAE generates the input embeddings. This VAE is trained to reconstruct a semantic latent space wherein similar FODs (in terms of peak count, profile variance, and peak directions) are mapped to nearby vectors, and dissimilar ones are mapped farther apart.

Inspired by the architecture introduced in[26–28], we utilized a basic VAE structure and a feedforward network to enhance explainability of the latent space. the encoder and decoder consist of *N*_*B*_ blocks, each composed of a linear layer, LayerNorm, and a SiLU activation. The decoder mirrors the encoder architecture. To normalize the SHCs, we normalize the SHCs using L2 norm. The feedforward network (shadow network) is composed of three linear layers followed by a softmax output layer.

Each input sample is denoted as *S*_*i*_ = {*SHC*_*i*_, *P*_*i*_}, where *SHC*_*i*_ is the spherical harmonic coefficient vector for the sample, and 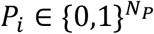 is a one-hot vector representing the number of FOD peaks (extracted via DIPY). Here, *N*_*P*_ is the maximum number of peaks. The encoder learns a Gaussian distribution *N*(*μ*_*i*_, *σ*_*i*_) from which the latent representation *Z*_*i*_ is sampled. The decoder reconstructs the estimated coefficients 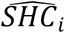 from *Z*_*i*_, while the feedforward network predicts 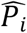.The training loss for the VAE is

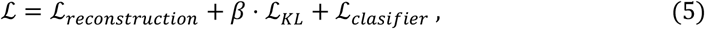

where *β* is a Kullback-Leibler (KL) divergence penalty and

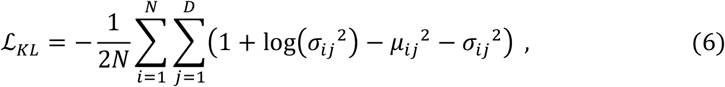

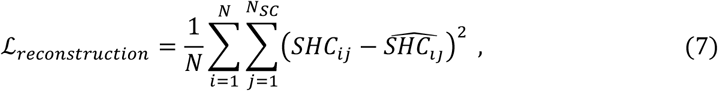

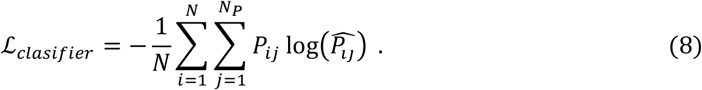

Here, *N* is the number of samples, and *D* is the latent dimension.

#### 2.2.2. Self-supervised learning strategy

The model is trained with a self-supervised objective by masking the central voxel’s SHCs during training. The network is optimized by minimizing the Mean Squared Error (MSE) between the FOD of the center voxel and the symmetric components of the estimated A-FOD [8]. The predicted parameter space embedding is used for reconstructing the estimated A-FOD. Notably, the A-FOD is represented by full basis SHCs [2, 4]. Subsequently, the symmetric portion of the estimated A-FOD is obtained by projecting only even order SHCs back onto the spherical domain [2, 8]. In essence, the proposed self-supervised Transformer model serves to define the shape of probability distributions governing A-FOD estimation. As a result, A-FODs will be aligned with the underlying fiber structures learned from data.

#### 2.2.3. Model Configuration

##### VAE Configuration

The VAE component of the architecture consists of three encoder and decoder blocks, *N*_*B*_ = 3, with layer sizes [128,64,32] and a latent dimension *D* = 10. The maximum number of peaks is set to *N*_*P*_ = 5. The KL divergence penalty in the loss function is set to *β* = 1. The model is trained using the Adam optimizer over 500 epochs with a batch size of 256 and a learning rate of 1e-3.

##### Transformer Configuration

The proposed Transformer model incorporates five Transformer Encoder blocks, each with a fixed latent dimensionality of D = 10. The MSA and MLP sub-blocks use the SiLU activation function. The feedforward head uses the layer sizes [4D,4D,5] to create parameter space embeddings. The Adam optimizer is used to train the model over 10 epochs with a batch size of 64 and an initial learning rate of 1e-3.

### 2.3. Material

#### The diffusion-simulated connectivity (DiSCo)

[14] is described in brief in the following. This numerical Phantom was designed to mimic complex anatomical fiber pathway trajectories. The streamlines correspond to the centerline trajectories of 12,196 synthetic tubular fibers, optimized using the Numerical Fiber Generator. To generate the corresponding dMRI signals, these trajectories were assigned gamma-distributed diameters (1.4 − 4.2 *μm*) and converted into 3D triangular meshes representing intra-axonal and myelin compartments. These meshes served as the substrate for Monte Carlo simulations of spin dynamics, ensuring the resulting signals are physically derived from the specific microstructure aligned with the pathways. The synthetic DW-MRI data was generated with an echo time (TE) of 53.5 ms and image dimensions of 40 × 40 × 40 voxels (1.0 mm isotropic). The acquisition scheme included four non-diffusion-weighted images and 360 diffusion-weighted directions distributed across four b-values (b = 1000, 1925, 3094, and 13,191 s/mm^2^).

#### *Multi-shell Human Dataset*[15]

The preprocessed dataset consists of volumes with an image dimensions of 146 × 146 × 92. For the purpose of this study, only the subject ses-c01r1 is included in the analyses. The diffusion scheme, containing 90 gradient directions (*b* ∈ {1000,2000,3000} s/mm^2^, colinear between any two shells) and 6 *b* = 0 s/mm^2^ volumes, was generated from a multi-shell vector sampling tool [29]. Preprocessing was carried out through the following pipeline. Denoising and Gibbs ringing correction[30] were performed using MRtrix3[31]. Motion and distortion correction were applied using tools provided in the FSL suite. Image distortion was estimated via the TOPUP tool, utilizing non-diffusion-weighted images with opposite phase-encoding directions to compute a field map [32]. Both distortion and motion were jointly corrected with EDDY tool[33]. Moreover, gradient directions were rotated accordingly to maintain alignment after EDDY-based corrections. All the analyses were performed within a mask obtained by applying a fractional anisotropy (FA)[34, 35] threshold of 0.25.

For both the DiSCo and Multi-shell Human Dataset, the symmetric FODs are reconstructed using multi-shell multi-tissue constrained spherical deconvolution (MSMT-CSD[36]). This method utilizes the information from all available b-shells to improve the reconstruction. The process is implemented in MRtrix3[37] with a maximum spherical harmonic (SH) order of 8.

### 2.4. Quantitative Metrics

To facilitate the quantitative evaluation of the generated A-FODs, we employ metrics that focus on two characteristics: directional asymmetry and deviation from standard symmetric representations.

#### Asymmetry index (ASI)

To measure the degree of asymmetry in the estimated A-FOD, we use the similarity measurement introduced in [4]. This measurement compares the A-FOD value along a direction *u* with the value along its opposite direction −*u*. This similarity is then used to compute the asymmetry index. The following expressions define the index:

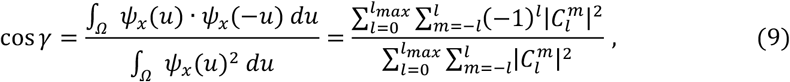

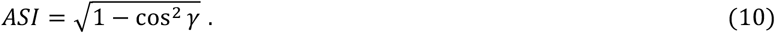

Here, 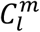 denotes the SH coefficient corresponding to the order *l* and degree *m* describing the estimated A-FOD profile, while cos *γ* represents the cosine similarity.

#### Model discrepancy index (MDI)

To quantify the variability between the asymmetric and symmetric FOD, we employ the model discrepancy index (MDI) described in [9], which is defined as

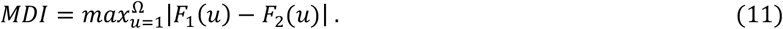

Here, Ω denotes the set of all directions on the unit sphere in the orientation distribution. *F*_1_ *and F*_2_represent the asymmetric and symmetric FODs, which are normalized to range [0, 1].

### 2.5. Performance Evaluation

The performance of the EnCAR method is evaluated through a combination of comparative and intrinsic analyses. The comparative analysis includes both qualitative visual inspection of the reconstructed A-FOD maps and a quantitative comparison using the ASI and MDI metrics. For fair comparison, we benchmark against two published methods operating within the general filtering framework described in Eq. (1):

- *Unified Filtering* [8]: This approach accounts four factors including inter-voxel distance, directional alignment, orientational similarity, and intensity similarity into a unified regularization weight. Each factor is modeled via a normal distribution, with optimal parameters values determined through a self-supervised grid search that minimizes the error between the input FOD and the symmetric component of the estimated A-FOD.
- *Tractosemas* [12]: This method estimates a field of asymmetric spherical functions from intra-voxel displacement probability. Its regularization weight is constructed from three factors: inter-voxel distance (modeled with a multivariate normal distribution) and both directional alignment and orientational similarity (modeled with von Mises–Fisher distributions).

All baseline methods are evaluated using their original parameter configurations. Where applicable, symmetric FODs from MSMT-CSD and ground-truth streamlines from the DiSCo phantom are included as additional references in the comparisons.

The intrinsic analysis involves generating maps of the individual learned regularization parameters to examine how the model adapts the shape of its weighting distributions in response to different local fiber geometries.

## 3. Results

### 3.1. DiSCo Phantom

We evaluated the performance of the EnCAR method on the DiSCo phantom where a cubic 3 × 3 × 3 kernel is used to define the neighborhood voxel set *N*(*x*).

#### 3.1.1. Intrinsic Analysis

The A-FOD map is illustrated in Fig. 3a from an axial slice (Z=24) of the DiSCo phantom, with glyphs reconstructed by EnCAR and normalized by their maximum amplitudes. In regions corresponding to single-fiber bundles, unidirectional patterns are observed, where glyphs exhibit symmetric shapes. In complex regions, the Y-shape, bending and uneven crossing patterns are revealed. The reconstructed A-FODs retain multiple peaks with clear angular separation. The lobes exhibit sharp, well-defined profiles, devoid of the blurring or over-smoothing that obscures directional information.

**Figure 3.**
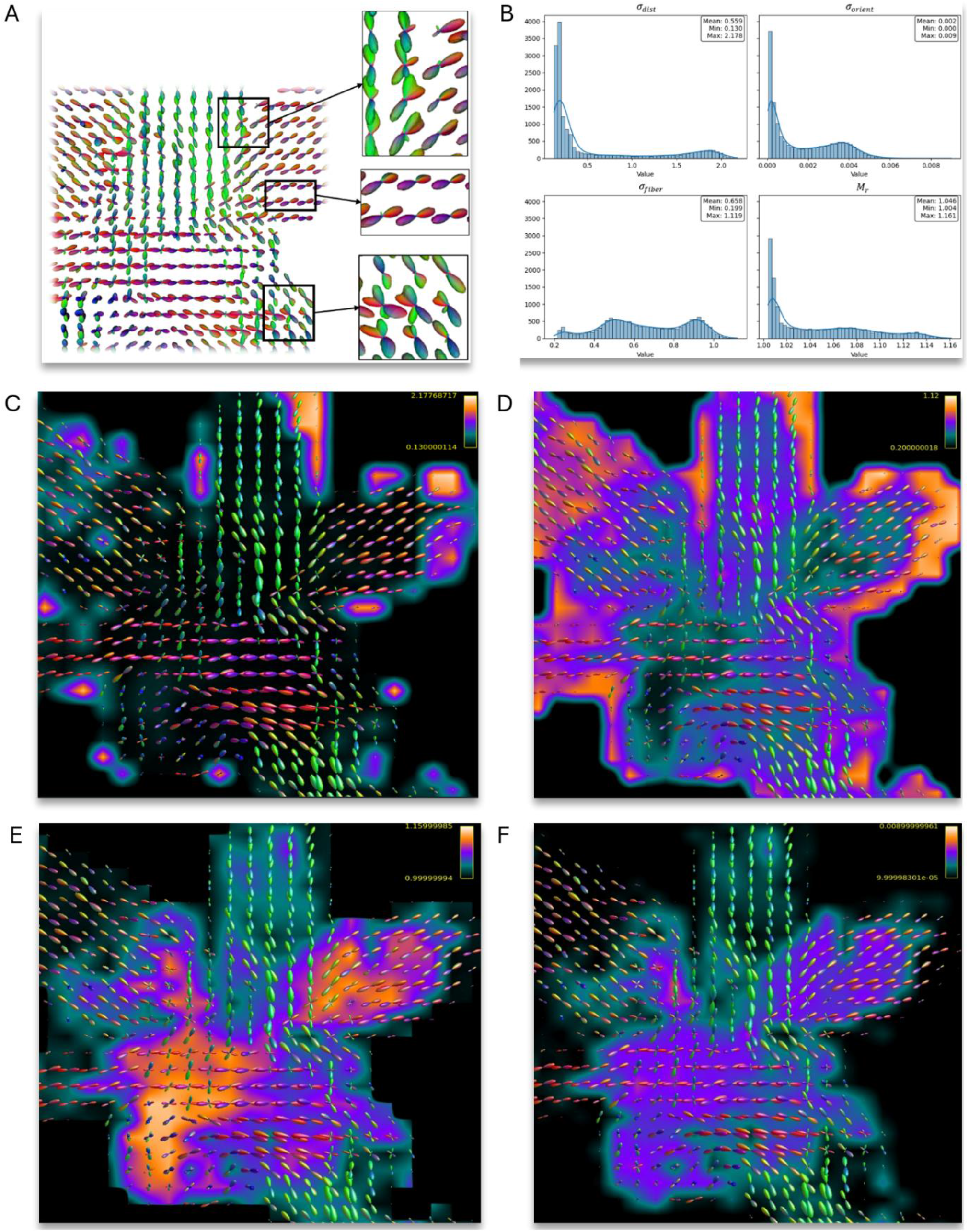
Intrinsic analysis on the DiSCo phantom. (A) The reconstructed A-FOD map for an axial slice (Z=24) (B) Histograms showing the distributions of the learned regularization parameters. (C-F) Voxel-wise heat maps of the individual parameters *(σ*_*dist*_, *σ*_*fiber*_, *M*_*r*_, *σ*_*orient*_ ), illustrating how the model adaptively tunes its behavior in response to local fiber geometry.

The voxel-wise heat maps of several regularization parameters (*σ*_*dist*_, *σ*_*fiber*_, *M*_*r*_ and *σ*_*orient*_) overlaid on the symmetric FODs are shown in Fig. 3c–3f. The spatial parameter *σ*_*dist*_ (Fig. 3c) displays higher values in boundary regions, reflecting the reduced reliability of FODs within *N*(*x*). Similarly, in these regions, *σ*_*fiber*_ (Fig. 3d) exhibits elevated values, increasing the contribution of available neighbors to the A-FOD reconstruction. Additionally, *σ*_*fiber*_ shows lower values in areas of low directional correlation between lobes compared to homogeneous regions.

The voxel-wise map of the regularization parameter *M*_*r*_, which modulates the maximum allowable rotation (Eq. 2) based on the inter voxel distance, is shown in Fig. 3e. The results indicate that non-zero rotation values are applied throughout the complex regions. Despite variation in *σ*_*orient*_ (Fig. 3f), analysis of its histogram (shown in Fig. 3b) reveals that the model consistently reduces this parameter to narrow range, with *σ*_*orient*_ ∈ [0.0001,0.009].

The histograms of the learned parameters across the phantom (Fig. 3b), summarize the model’s behavior. The *σ*_*dist*_ ∈ [0.13,2.18] shows that EnCAR utilizes FODs with non-uniform influences. The distribution of *M*_*r*_ ∈ [1.00,1.16] shows an average rotation of 0.046 rad (≈ 2.63°) at the corner of *N*(*x*), with a subset of voxels (where *M*_*r*_ = 1.00) corresponding to straight-path assumption. For directional alignment, the model utilizes *σ*_*fiber*_ ∈ [0.20,1.12], with a mean of 0.66.

#### 3.1.2. Qualitative Visual Inspection

To visually assess reconstructed glyphs, Fig. 4a-4c compares the A-FODs estimated by EnCAR against the two A-FOD methods, Tractosemas and Unified Filtering, and the symmetric MSMT-CSD FODs. Streamlines representing the midline of the axon 3D trajectories are overlaid on each map to serve as a reference for alignment accuracy.

**Figure 4.**
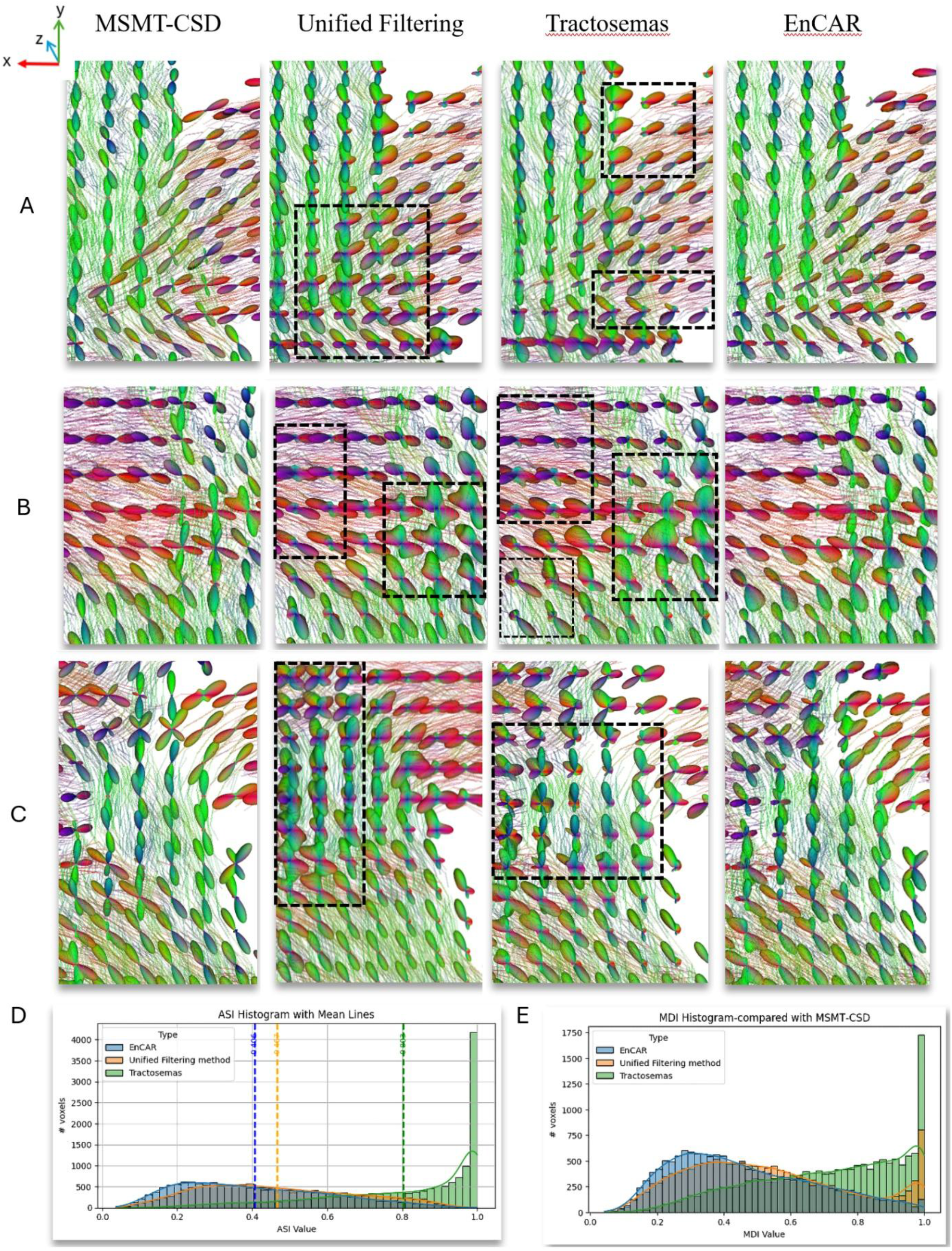
Qualitative and quantitative evaluation of A-FOD reconstruction methods on the DiSCo phantom. (A–C) Qualitative comparison in three distinct regions (one per row). The A-FODs generated by the EnCAR method are visually compared against two baseline methods (Unified Filtering and Tractosemas) and the initial MSMT-CSD results. To validate the alignment with local fiber geometry, the streamlines are overlaid on each map. (D) Asymmetry Index (ASI) distributions across the evaluated methods. (E) Model Discrepancy Index (MDI) distributions, assessing deviation from the reference FODs derived by MSMT-CSD.

In the region depicted in Fig. 4a, the Tractosemas method preserves the primary lobes of the A-FOD glyphs but shows signal voids (incomplete glyph structures) along boundary regions and within complex areas characterized by bending and fiber crossings. While Unified Filtering produces a more continuous field than Tractosemas, the glyphs in crossing regions appear diffuse with fused lobes (high profile variance). In contrast, EnCAR displays continuous A-FODs with distinct lobes that follow the streamline trajectories.

Both Tractosemas and Unified Filtering methods introduce artifacts in the region shown in Fig. 4b, estimating spurious peaks in areas that contain only unidirectional fibers. Both approaches are unable to resolve distinct lobe structures within the crossing regions and the glyphs appear with high profile variance. Conversely, the EnCAR method estimates A-FODs with sharp, high-intensity lobes that are structurally consistent with the local fiber pathway. It reconstructs distinct Y-shaped and bending A-FOD configurations without the spurious peaks observed in the baseline methods.

The Unified Filtering method generates valid A-FODs in only a subset of the unidirectional fiber regions (Fig. 4c). It exhibits glyphs characterized by high variance lobes. In these regions, Tractosemas yields incomplete glyph structures and discontinuous field. In contrast, the proposed method consistently presents high-intensity lobes aligned with the curvature of local fiber bundles.

#### 3.1.3. Quantitative Comparison

The quantitative evaluation using histograms of the ASI and MDI metrics is summarized in Figs. 4d and 4e. The ASI distributions are generally similar for EnCAR and Unified Filtering. The Tractosemas distribution, however, exhibits higher overall ASI values. To verify that EnCAR’s ASI distribution is robust, a variance analysis is provided in Appendix A.

Regarding the MDI distributions (Fig. 4e), where MSMT-CSD serves as the reference, the EnCAR method displays a unimodal distribution skewed toward lower values. In contrast, the MDI histogram for the Unified Filtering method features a distinct secondary peak at MDI = 1, while Tractosemas shows a predominant peak at MDI = 1.

### 3.2. Multi-shell Human Dataset experiments

To evaluate performance in real-world scenarios, we applied our method to the in-vivo multi-shell human dataset. For this analysis, the neighborhood voxel set *N*(*x*) is defined using a cubic 5 × 5 × 5 kernel, allowing us to effectively demonstrate the adaptive behavior of the regularization parameter *M*_*r*_.

#### 3.2.1. Intrinsic Analysis

Histograms of the learned regularization parameters are presented in Fig. 5. The parameter *σ*_*fiber*_ ∈ [0.15,1.34] exhibits a broad distribution, varying across the voxels. The distribution of *M*_*r*_ ∈ [1.23,1.40] shows a minimum rotational correction of 0.23 rad (≈ 13.15°) at the corner of *N*(*x*). Regarding the spatial parameter *σ*_*dist*_, the distribution shows that only direct neighbors (*I*_*xy*_= 1) within *N*(*x*) are utilized to reconstruct the A-FODs. Furthermore, *σ*_*orient*_ converges to 0.002.

**Figure 5.**
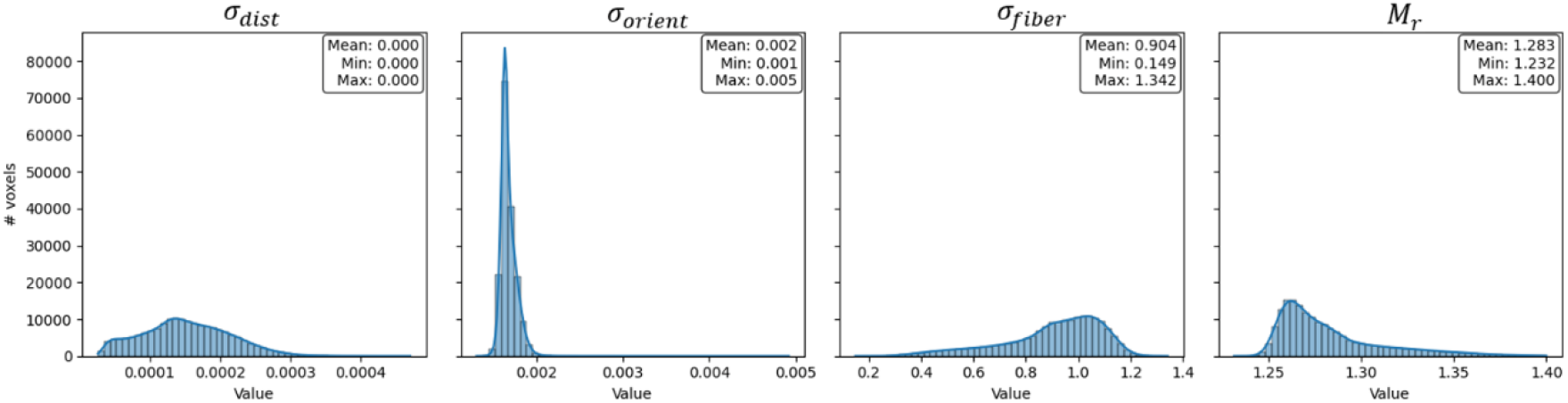
Histograms of the learned regularization parameters for the multi-shell human dataset.

The heat maps of the learned parameters *M*_*r*_ *and σ*_*fiber*_ are presented in Fig. 6 for two axial slices (Z=45 and Z=59), overlaid on the symmetric FODs. Regarding *σ*_*fiber*_, elevated values are observed in regions where increasing the contribution of neighboring voxels is beneficial. This occurs near the boundaries of the brain mask, where the neighborhood *N*(*x*) is incomplete and not all voxel information is available, and in regions exhibiting high directional correlation between FOD lobes. Conversely, *σ*_*fiber*_ values are reduced in complex zones.

**Figure 6.**
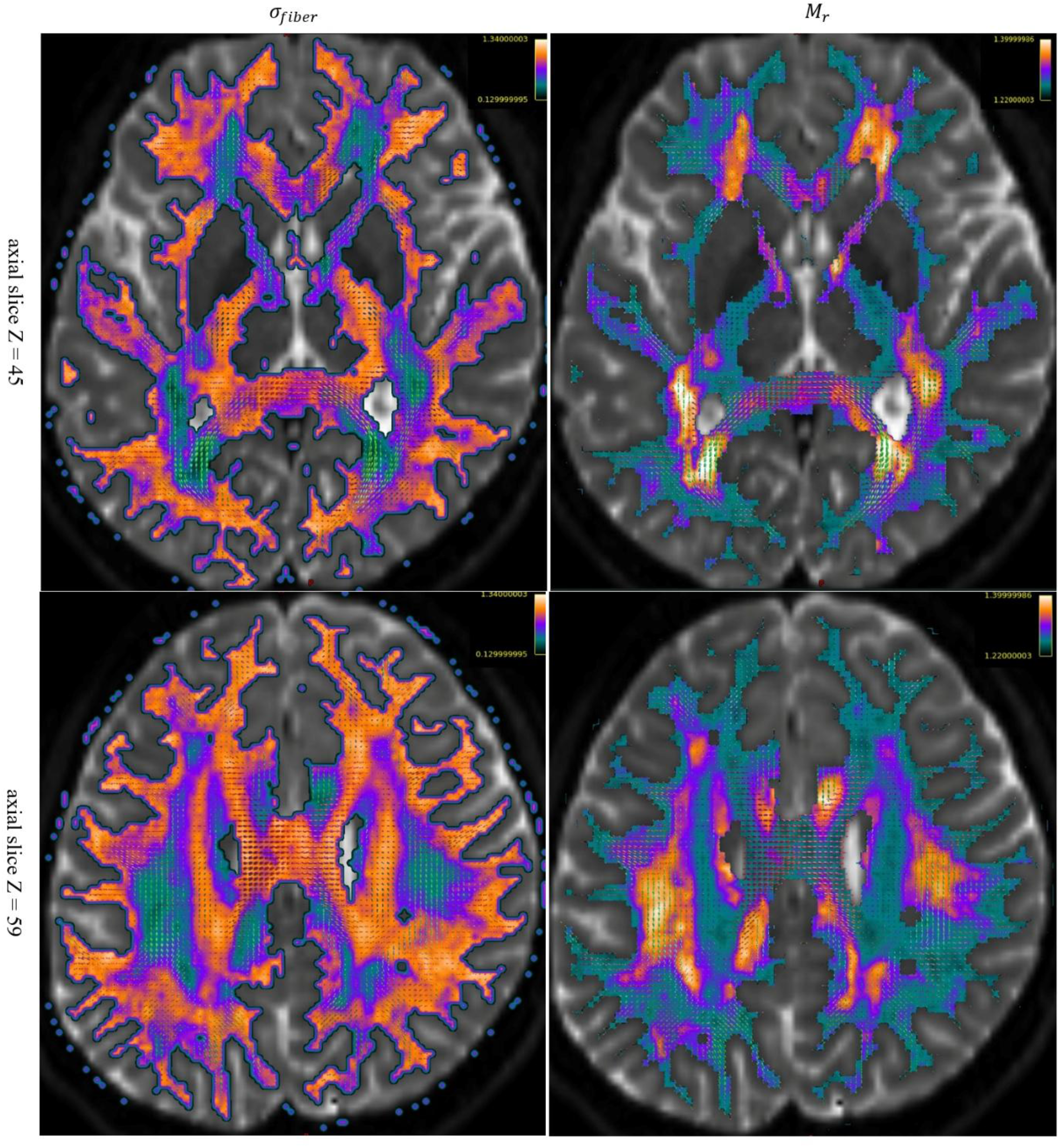
Intrinsic analysis on the multi-shell human dataset. Voxel-wise heat of the *σ*_*dist*_ (left column) and *M*_*r*_ (right column) shown for two axial slices (Z=45 and Z=59). These maps illustrate how the model adaptively tunes its behavior in response to the local fiber geometry.

Regarding *M*_*r*_ (rotational corrections), the maps exhibit low values in regions where neighboring FODs are highly correlated and have similar orientations. In contrast, in regions of high angular complexity and lower FOD correlation, *M*_*r*_ values are elevated. Additionally, *M*_*r*_ remains low in peripheral regions.

#### 3.2.2. Qualitative Visual Inspection

Qualitative comparison of A-FOD glyphs in selected regions of an axial slice (z = 45) from the in vivo multi-shell dataset is shown in Fig. 7a. The aim of this comparison is to assess each method’s performance on tract continuity, angular resolution, and orientation accuracy.

**Figure 7.**
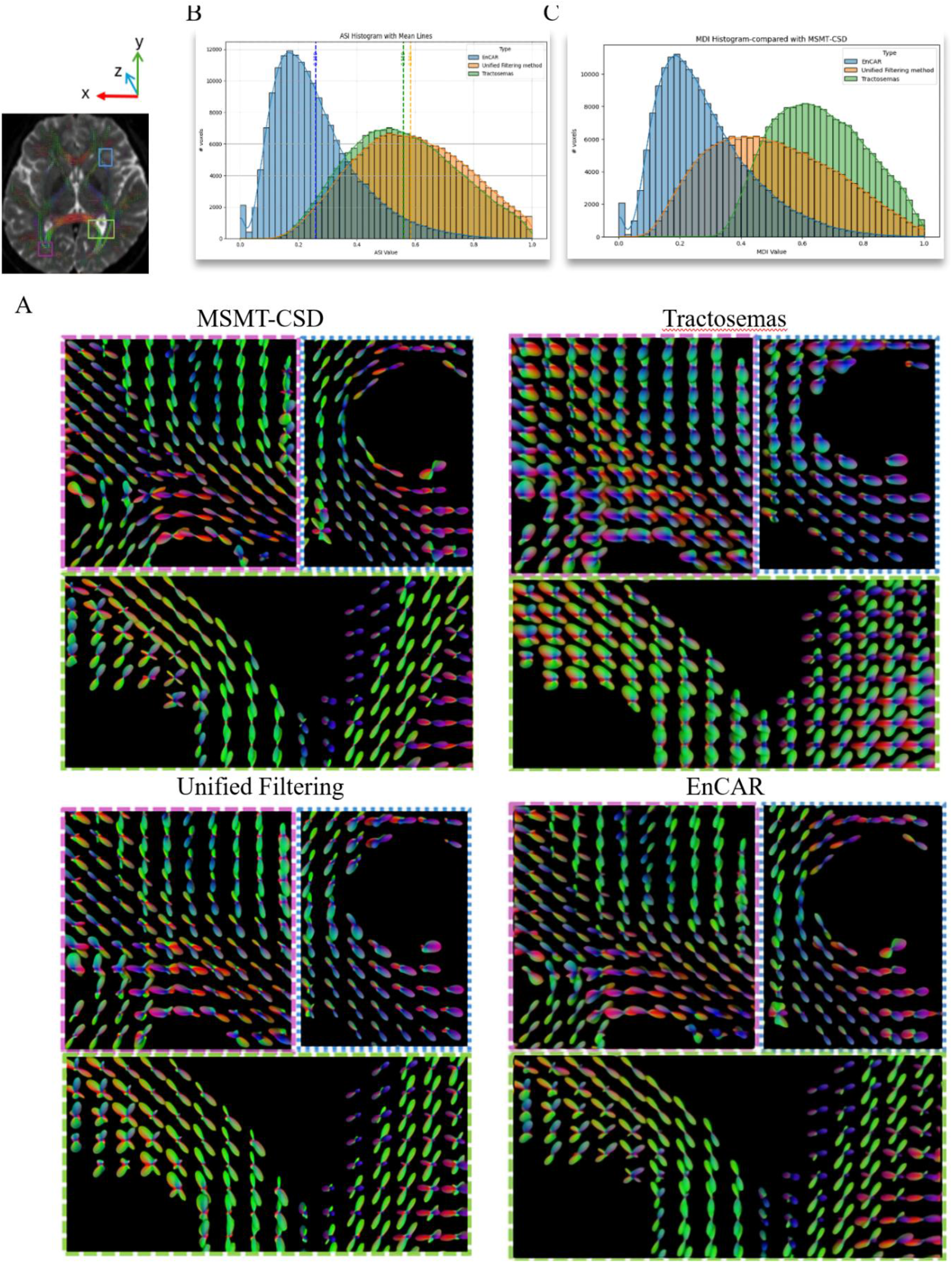
Qualitative and quantitative comparison of A-FOD reconstruction methods on the multi-shell human dataset. (A) Qualitative comparison in selected regions of an axial slice (Z=45), the A-FODs generated by the EnCAR method are compared against the baseline approaches (Unified Filtering and Tractosemas), with the initial MSMT-CSD FODs included as a reference for the underlying fiber orientations. (B) Asymmetry Index (ASI) distributions across the evaluated methods. (C) Model Discrepancy Index (MDI) distributions, assessing deviation from the reference FODs derived by MSMT-CSD.

In regions near anatomical boundaries or where neighborhood information (FODs with in *N*(*x*)) is limited, both Tractosemas and Unified Filtering produce incomplete A-FODs and exhibit abrupt variations in glyph transitions across voxels. In the area with lower correlation between neighboring lobes’ orientations, the glyphs produced by baseline methods appear more isotropic compared to EnCAR. Specifically, Tractosemas generates A-FODs with more isotropic profiles (lower angular resolution) compared to Unified Filtering. In addition, comparing them to the MSMT-CSD reference shows that the orientations encoded by the baseline methods are frequently misaligned with the underlying fiber geometry.

In contrast, the EnCAR method reconstructs continuous A-FOD maps even in boundary zones. The estimated glyphs exhibit sharp, high-angular-resolution profiles with distinct lobes. The orientation of these lobes maintains alignment with the MSMT-CSD reference, even in regions characterized by Y-shaped, bended and uneven A-FOD configurations.

#### 3.2.3. Quantitative Comparison

The histograms of the ASI and MDI metrics for the in-vivo dataset are presented in Fig. 7b and 7c. In the ASI analysis, both Tractosemas and the Unified Filtering method exhibit similar distributions, with mean ASI values of 0.55 and 0.58, respectively, and a notable accumulation of voxels at the upper bound (ASI = 1). In contrast, EnCAR demonstrates a distinct profile characterized by a left-skewed distribution (mean ASI = 0.25) and a subset of voxels at the lower bound (ASI=0). The variance analysis of ASI is provided in Appendix A.

In the MDI analysis (Fig. 7c), the distribution for EnCAR is shifted toward lower values relative to both baseline approaches and the reconstructed A-FODs exhibit the lowest discrepancy relative to the symmetric MSMT-CSD reference. Conversely, the baselines exhibit distributions centered at higher values, with a population of voxels at the maximum discrepancy (MDI = 1).

## 4. Discussion

By employing a self-supervised paradigm, EnCAR resolves sharp, high-angular-resolution A-FODs in regions characterized by bending, branching, and fanning. Specifically, it reconstructs T- or Y-shaped and uneven crossing patterns, as demonstrated in both the DiSCo phantom and the in vivo multi-shell human dataset.

### 4.1. Overcoming Straight-Line Constraints

A central limitation of existing A-FOD filtering methods [4, 8, 10, 12, 13] is the simplifying assumption that fibers propagate linearly between the central voxel and all neighboring voxels within *N*(*x*). Results demonstrate that this assumption causes baseline methods to aggregate unrelated directional information in complex and curved regions, resulting in the isotropic and blurred glyphs lacking clearly separable lobes, as observed in Fig. 4 and Fig. 7. These methods, by restricting information modeling to straight trajectories, have insufficient capacity to accurately represent curved fiber paths. Although, the radius of curvature of white matter fibers is typically larger than individual voxel dimensions[38], curved fiber paths still occur over multiple adjacent voxels in the neighborhoods where the straight-line assumption fails to capture the local fiber geometry. The EnCAR method overcomes this limitation via a learned rotational transformation governed by *S*_*p*_ and *M*_*r*_. This mechanism provides the model with the necessary degrees of freedom to adaptively utilize curvature features, thereby enforcing better continuity with its neighbors along the estimated fiber segment. The necessity of this correction is substantiated by the fact that the model applies non-zero rotations selectively (Fig. 3e and Fig. 6). While *M*_*r*_ = 1 in aligned regions, the mean correction of *M*_*r*_ > 1 confirms that the straight-path assumption along *U* and −*U* fails in complex regions, particularly in regions exhibiting low angular correlation between voxels [39]. By rotating orientation vectors, EnCAR incorporates geometrically meaningful information rather than conflicting signals.

Consequently, we achieve high-angular-resolution glyphs even in areas where baseline methods fail to resolve dominant fiber directions, thereby facilitating the reconstruction of smoother fiber segments.

### 4.2. Region-specific regularization parameters

The reliance on a fixed set of global regularization parameters is addressed by our self-supervised Transformer, which dynamically adjusts the regularization parameters (*σ*_*dist*_, *σ*_*fiber*_, *σ*_*orient*_, *S*_*p*_ and *M*_*r*_) in response to local fiber geometry. Functioning as a local adaptive probability kernel, this parameter set governs the relative influence of the surrounding FODs during the estimation process. Given the diverse geometric configurations present across the voxels [7, 40, 41], imposing a stationary global probability kernel is inadequate, leading to the aggregation of incoherent directional signals in complex regions. In contrast, EnCAR adapts its weighting strategy based on regional complexity. As evidenced by both the DiSCo phantom (Fig. 3c) and the human dataset (Fig. 6), the model learns to decrease *σ*_*fiber*_ in complex crossings and curved regions to preserve angular detail. Conversely, in boundary regions with sparse FODs and areas of highly correlated FODs, the model increases *σ*_*fiber*_ to incorporate sufficient neighborhood information. The dynamic behavior across varying levels of angular complexity validates the use of learnable kernels to govern the A-FOD estimation process.

The performance of Transformer-based architectures is dependent on the representational quality of their input features[42–44]. Raw SH coefficients, while theoretically complete descriptors of the fiber orientation distribution function, suffer from several well-known limitations in practice: high dimensionality, rotational ambiguity in their representation[45–47], and a lack of explicit semantic organization with respect to fiber geometry or directional continuity. These characteristics impose a substantial burden on the model, forcing it to simultaneously learn both low-level decomposition and high-level reasoning from high-dimensional inputs. Our proposed SH Semantic Encoder addresses this challenge by projecting the raw SH coefficients into a compact, highly structured latent space that respects both directional similarity and geometric complexity (Appendix B, Fig. B.1). The latent space supports linear arithmetic operations that correspond to interpretable semantic changes in the diffusion profile, properties that are largely absent in the original SH coefficient space (Appendix B, Fig. B.2). By supplying the Transformer with these semantically rich, low-dimensional embeddings rather than raw coefficients, we simplify the learning objective.

### 4.3. Quantitative Fidelity and Implications

Quantitative evaluation confirms that EnCAR yields significant improvements over Unified Filtering [8] and Tractosemas [12]. First, the MDI analysis (Fig. 4e, Fig. 7c) reveals that our reconstructed A-FODs exhibit the lowest discrepancy relative to the symmetric MSMT-CSD reference [9]. This suggests that the proposed method enhances structural information, via the asymmetric component, without deviating from valid diffusion properties. The baseline methods exhibit a subset of voxels with MDI = 1 where the reconstructed A-FODs are entirely dissimilar to the reference symmetric FODs, indicating a severe loss of structural coherence in challenging regions. Second, the left-skewed ASI distribution (Fig. 7b) observed in the human dataset offers a key insight into the model’s behavior. Unlike baseline methods, which exhibit a global, right-skewed asymmetry distribution, our model displays a peak at low ASI values and a subset of voxels at the lower bound (ASI=0). This suggests an adaptive mechanism that preserves symmetry in regions where the underlying fiber paths are expected to be symmetric and applies asymmetry only where geometrically necessary[4, 9]. The baselines exhibit elevated ASI values. Our qualitative analysis suggests that this should not be interpreted as superior resolution of complex structures. Rather, the high ASI values, particularly on voxels with ASI=1, are attributable to the frequent generation of incomplete or truncated A-FODs, which artifactually inflate the asymmetry index without representing true anatomical features such as fiber curvature, bending, or fanning. Thus, the EnCAR method provides a more meaningful A-FOD representation.

The accurate estimation of sharp A-FODs aligned with the underlying structure results in more coherent and smooth directional transitions across voxels [2]. These smooth transitions provide a way to gain a greater understanding of trends over a spatial neighborhood [8] and enhance the visibility of continuous fiber segments, which is expected to directly contribute to more reliable tractography [9, 13, 48, 49]. In contrast, the high-variance, ambiguous A-FODs produced by baseline methods obscure the separation of distinct fiber populations. This directional uncertainty impedes the independent tracing of intersecting bundles and frequently leads to premature streamline termination due to accumulated propagation errors in complex regions [50, 51].

### 4.4. Limitations and Future work

Despite its strengths, EnCAR performance is compromised in regions with insufficient or unreliable neighborhood information, such as tissue boundaries and mask edges, due to its inherent dependence on neighboring FODs within *N*(*x*). Although including the center voxel’s FOD would theoretically improve A-FOD reconstruction [4, 8], as this local signal is highly reliable, it must be excluded to define a valid self-supervised objective (Appendix C). More fundamentally, all A-FOD estimation methods, including the present work, lack true ground truth for A-FODs. Validation therefore relies on qualitative assessments like glyph inspection and visual alignment, complemented by quantitative metrics (ASI and MDI) that provide relative comparisons.

In future work, we intend to investigate architectures that enable the direct estimation of A-FODs from raw diffusion-weighted signals, thereby bypassing the intermediate symmetric FOD computation step. This approach has the potential to reduce error propagation and further enhance angular resolution. Furthermore, to explicitly validate the clinical utility of this method, we plan to conduct detailed tractography assessments quantifying the improvement in streamline connectivity for complex bundles.

## 5. Conclusion

We proposed the encoder-based curvature-aware regularization method for estimating asymmetric fiber orientation distributions. The method excludes the central voxel from the estimation process and uses a self-supervised Transformer with semantic spherical harmonics encoding to learn curvature-adjusted relationships between neighboring voxels. By learning the regularization parameters aligned with local fiber pathway, the method effectively models diverse fiber geometries across the brain. Experiments on DiSCo and Multi-shell Human Dataset showed that it consistently reconstructed high-angular-resolution A-FODs, maintained tract continuity, and accurately represented complex fiber shapes such as crossed, fan-shaped, branching and curved pathways. By adapting the regularization shape, the model maintains angular detail in challenging regions. It also ensured smooth continuity along coherent fiber bundles. Quantitative analyses verified that the method preserved the underlying diffusion signal more accurately.

## CRediT authorship contribution statement

M.T.: Writing – original draft, Methodology, Investigation, Conceptualization. M.P.: Writing – review & editing, Validation. M.M: Writing – review & editing, Validation. T.B.D.: Writing – review & editing, Validation, Investigation, Funding acquisition, Conceptualization.

## Data Availability

The datasets analyzed in this study are publicly available from their original sources. The DiSCo phantom [14] is openly accessible via Mendeley Data (https://data.mendeley.com/datasets/fgf86jdfg6/3). The Multi-shell Human Dataset [15] is accessible via the public repositories linked in the original publication (https://doi.org/10.6084/m9.figshare.8851955)

## Code Availability

The source code and trained models supporting the findings of this study will be publicly available on GitHub (https://github.com/MaP-science/EncoderBasedRegularization-AFOD) upon acceptance of the manuscript.

## Acknowledgments

M.T. was supported by the Smart Computing program, a joint Ph.D. program co-financed by the Universities of Florence, Pisa, and Siena. This project received funding from the European Research Council (ERC) under the European Union’s Horizon Europe research and innovation programme (Grant Agreement No. 101044180; Principal Investigator: Tim B. Dyrby). Furthermore, M.P. acknowledges funding from the Independent Research Fund Denmark (DFF; Case No. 10.46540/3105-00129B; Principal Investigator: Marco Pizzolato).

## Appendix A: Characterization of Model Stability

To evaluate the stability of the EnCAR, we assessed its performance on the DiSCo phantom and the in-vivo multi-shell human dataset. Specifically, the network was trained from scratch in six independent runs, utilizing random weight initializations for each iteration.

Fig. A.1a illustrates the histogram of the voxel-wise standard deviation (std) of the ASI for the DiSCo phantom over six independent runs, alongside an inset detailing the estimated A-FODs for three randomly selected voxels with *std* ≈ 0.15. The low average std value of 0.067 demonstrates that EnCAR exhibits high training stability and is robust to parameter initialization, confirming that the model converges to similar A-FODs across different runs. Regarding the analysis of the distribution’s tail, A mosaic in axial view of the voxel-wise std is shown in Fig. A.1b.

**Figure A.1.**
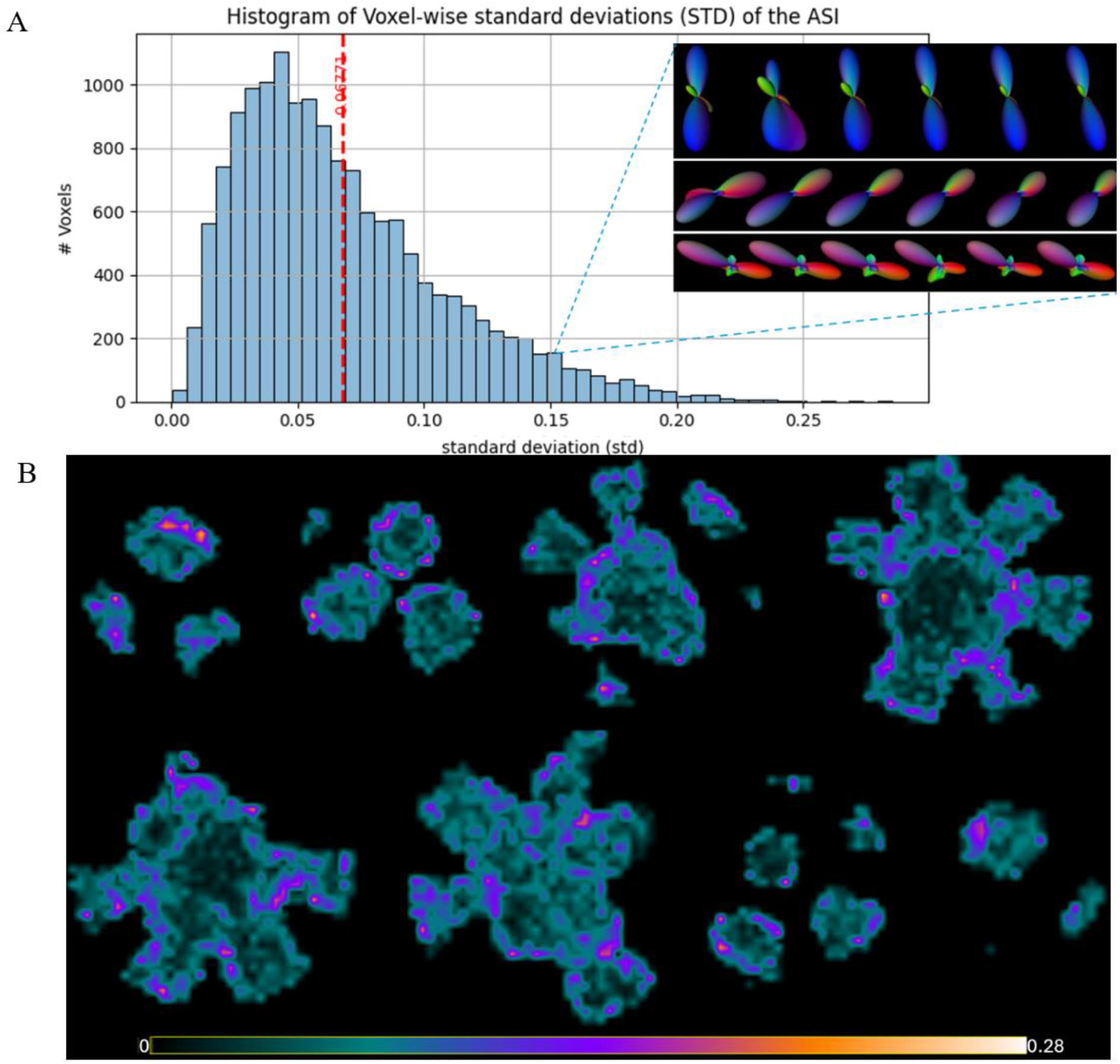
Evaluation of EnCAR stability across six independent runs in the DiSCo phantom. (A) Histogram of the voxel-wise standard deviation (std) of the ASI, alongside an inset detailing the estimated A-FODs of three random selected voxels (*std* ≈ 0.15). (B) Mosaic of the voxel-wise std (Axial view) of the ASI, visualizing the spatial distribution of variance across the phantom.

Spatial analysis reveals that the std peaks at the mask boundaries, where unreliable contextual information within *N*(*x*) drives up model uncertainty. Conversely, this variance progressively decreases toward the internal regions, as the information within *N*(*x*) becomes fully available and reliable.

A comparable stability profile is observed when applying the EnCAR to the multi-shell human dataset (Fig. A.2). The std histogram of ASI (Fig. A.2a) yields an even lower average std of 0.044, further confirming the model’s robustness to weight initialization in complex, real-world anatomical data. As shown in Fig. A.2b, the SD remains consistently low throughout the internal white matter regions, where reliable neighborhood information guides the self-supervised reconstruction. In contrast, model uncertainty is elevated at the peripheral boundaries.

**Figure A.2.**
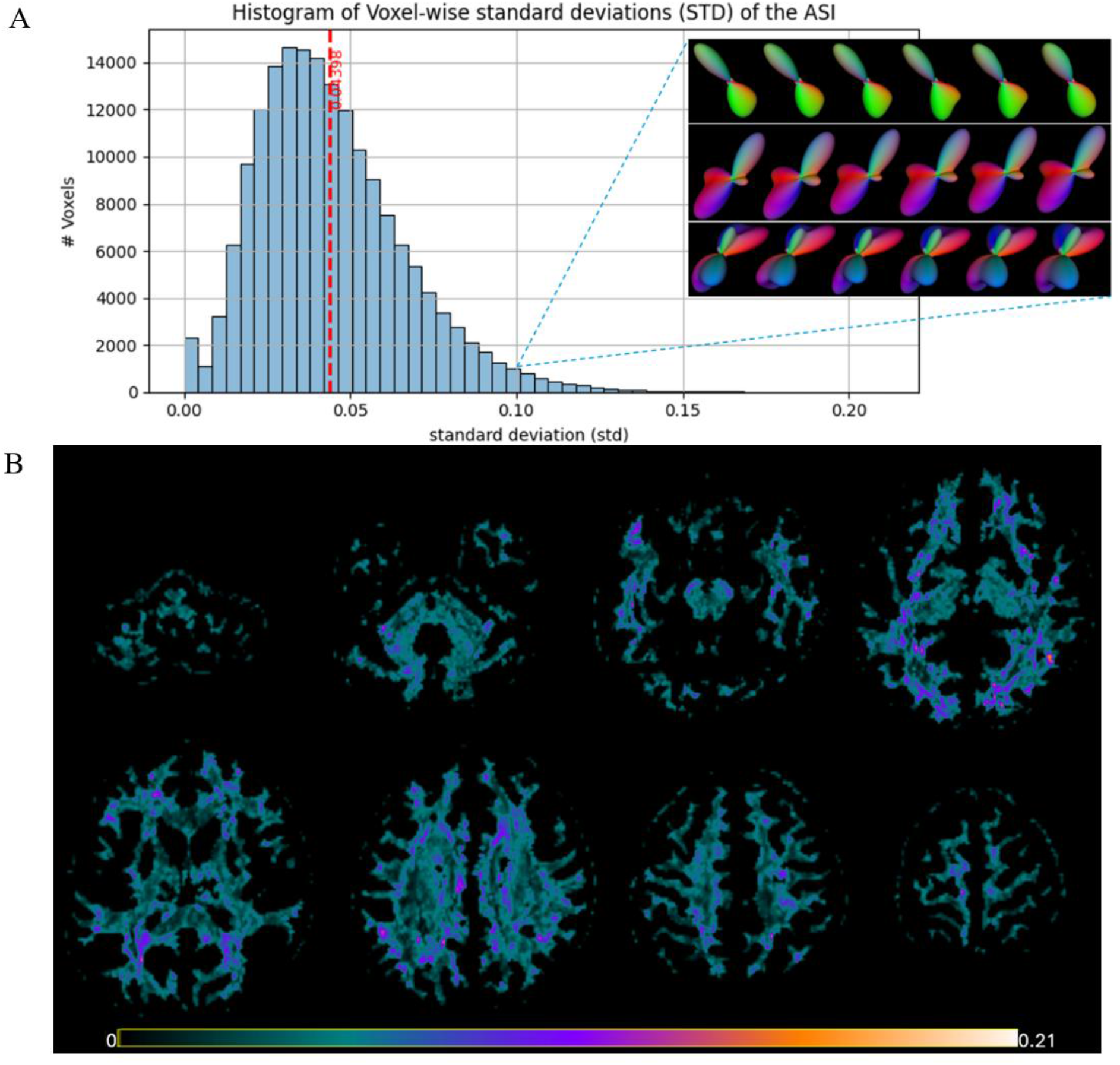
Evaluation of EnCAR stability across six independent runs in the multi-shell human dataset. (A) Histogram of the voxel-wise standard deviation (std) of the ASI, alongside an inset detailing the estimated A-FODs of three random selected voxels (*std* ≈ 0.10). (B) Axial mosaic of the voxel-wise std of the ASI, visualizing the spatial distribution of variance across the brain volume.

## Appendix B: Semantic Latent Space Characterization

Understanding the latent spaces learned by the VAE is important for interpreting how the self-supervised transformer reconstructs A-FODs. The encoder in the VAE produces input embeddings. These compact semantic embeddings reduce the complexity and number of parameters needed by the Transformer.

To gain insights into latent space, we use t-distributed Stochastic Neighbor Embedding (t-SNE) [52] to project the latent vectors into a two-dimensional Euclidean space for visualization. The t-SNE algorithm is used with default settings and perplexity value of 40. The latent space is visualized across all voxels from both datasets in Fig. B.1, where the color bar indicates the number of FOD peaks per voxel, ranging from 1 to 4.

The latent space visualization shows how the network differentiates FODs according to their directional complexity. Single-peak FODs tend to cluster along the outer periphery of the space, as indicated by the black points. Each outer region corresponds to a dominant diffusion direction, as shown in the decoded FODs. As the number of peaks increases, there is a clear transition toward the central region of the latent space, where more complex, multi-peak FODs appear, indicated by the red and yellow points. Multi-peak FODs are composed of several unidirectional components. As a result, they are positioned near their respective unidirectional components due to shared structures and peak orientations. This spatial organization demonstrates that the VAE has learned a structured representation that encodes a natural hierarchy based on FOD complexity. Beyond this overall structure, the latent space exhibits two key properties that confirm its well-ordered nature: locality and support for semantic arithmetic.

### Locality

The space’s locality was assessed by introducing small Gaussian noise to a latent vector and decoding the resulting FODs. Fig. B.2a shows two examples: in each case, the original decoded FOD is shown (left), and the right panels display multiple decoded FODs generated by adding noise to the same latent vector. This local neighborhood exploration reveals that the surrounding area structurally and directionally are similar to a selected FOD and minor changes in latent space produce smoothly varying FODs with consistent orientation and shape. This confirms that semantically similar FODs lie close together, and that the latent space exhibits strong locality properties.

### Semantic Arithmetic

In Fig. B.2b, we demonstrate an arithmetic operation in the latent space to assess how the model understands the meaning of FOD directions. To explore the semantic arithmetic capacity of the latent space, we perform vector operations involving multi-peak FODs. Specifically, we subtract the latent vector of an unwanted direction (FOD B) from a source (FOD A), and add another direction (FOD C), then decode the result. The result FOD reflects the intended directional transformation. This behavior confirms that the latent space supports linear arithmetic operations that correspond to interpretable semantic changes in the diffusion profile.

**Figure B.1.**
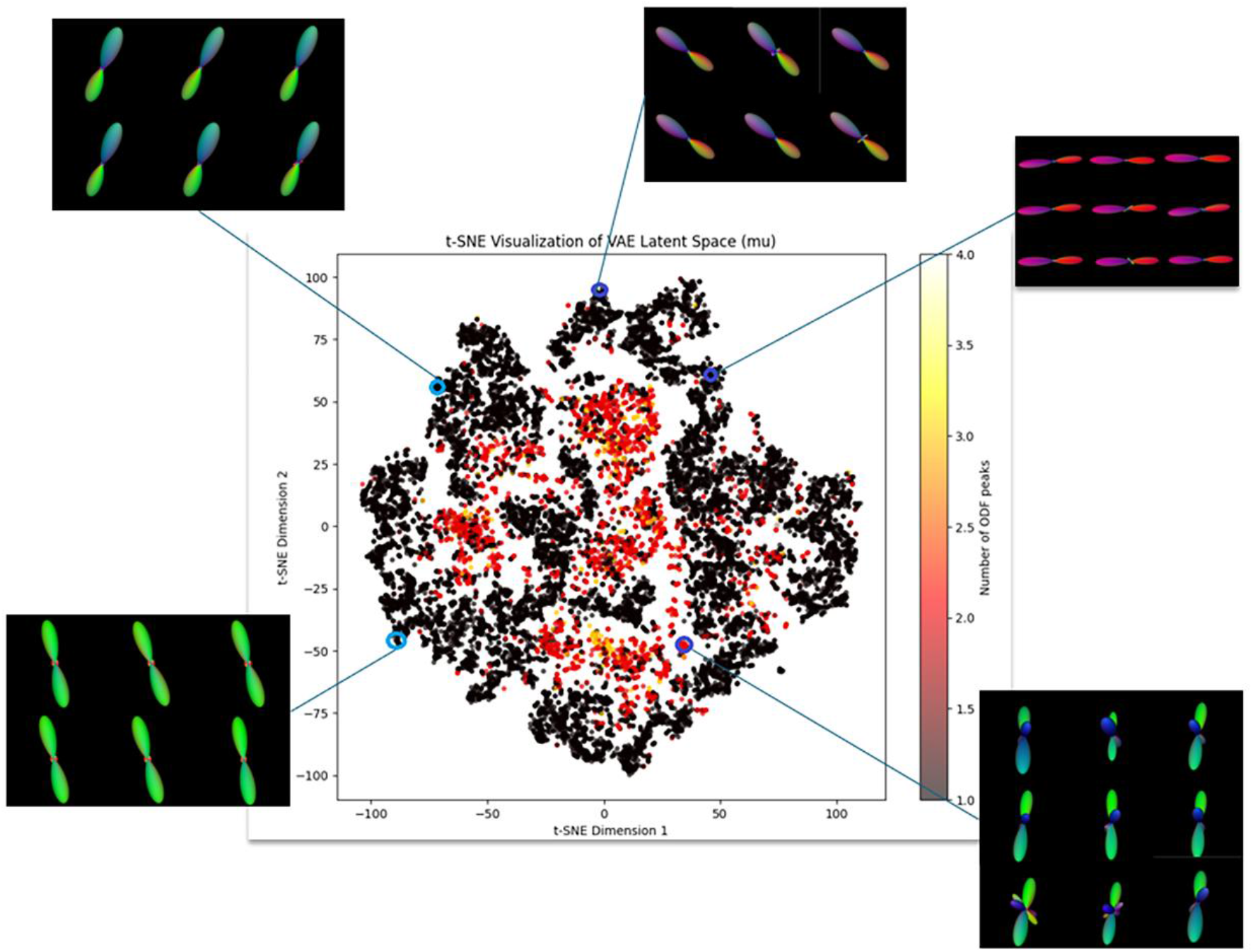
A t-SNE visualization of the VAE latent space, colored by the number of FOD peaks per voxel. The plot demonstrates a learned hierarchy based on complexity, with single-peak FODs mapped to the periphery and multi-peak FODs concentrated in the center.

**Figure B.2.**
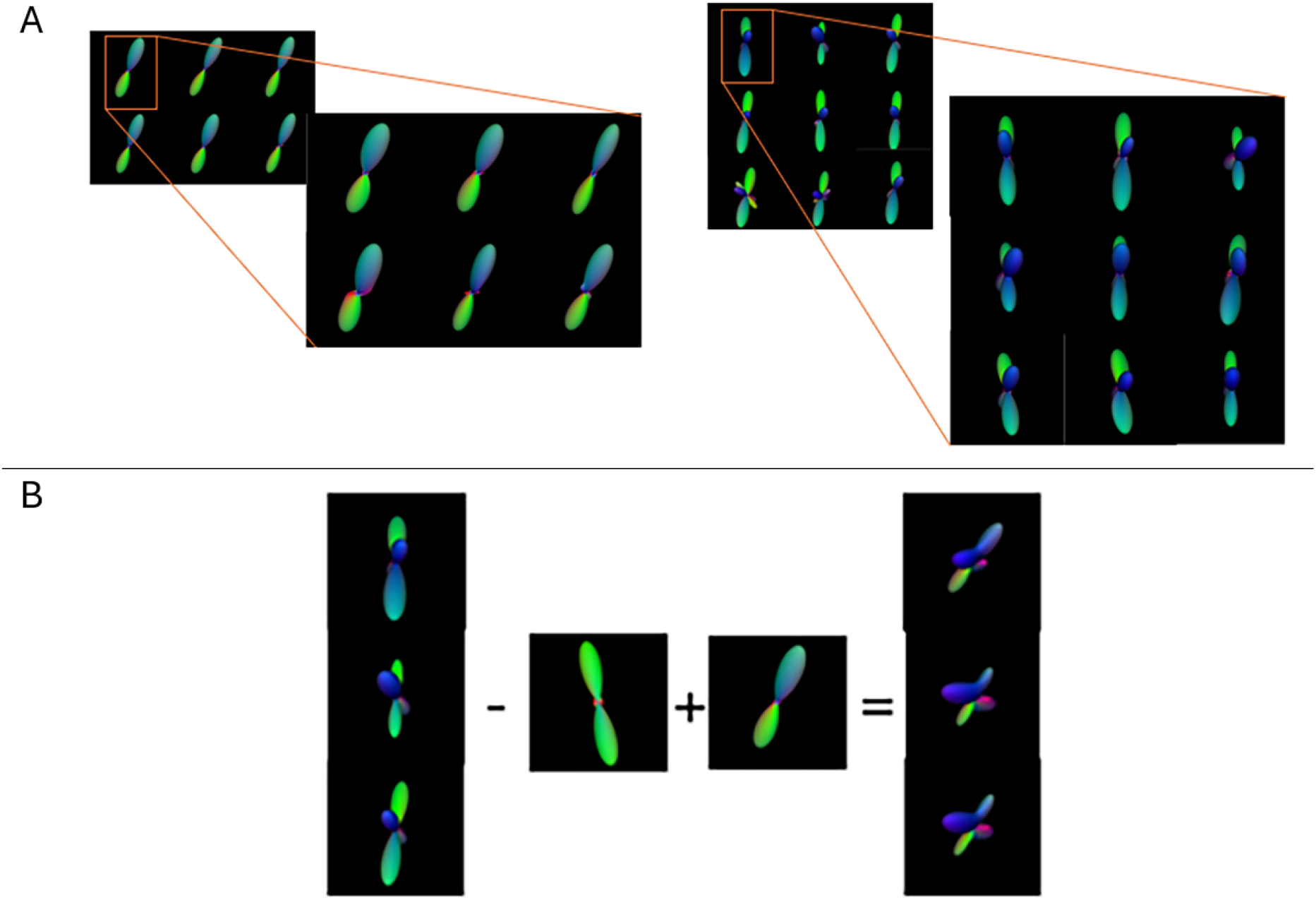
Characterization of the learned VAE latent space. (A) Demonstration of the locality property, where small perturbations of a single latent vector (left) result in decoded FODs (right). (B) Demonstration of semantic arithmetic, where operations on latent vectors produce a decoded FOD that reflects the intended semantic transformation.

## Appendix C: A-FOD Reconstruction Artifacts

Acknowledging the model’s limitation is essential for interpreting how EnCAR reconstructs A-FODs. To enable a valid self-supervised loss function during training the model, we explicitly exclude the center voxel’s FOD from *N*(*x*). As a result, the network estimates the central A-FOD using information solely from its spatial neighbors. This design choice can lead to blurred glyphs with poorly defined lobes in regions where reliable contextual information within *N*(*x*) is insufficient. Fig. C.1 provides representative examples of these artifacts in the DiSCo phantom.

**Figure C.1.**
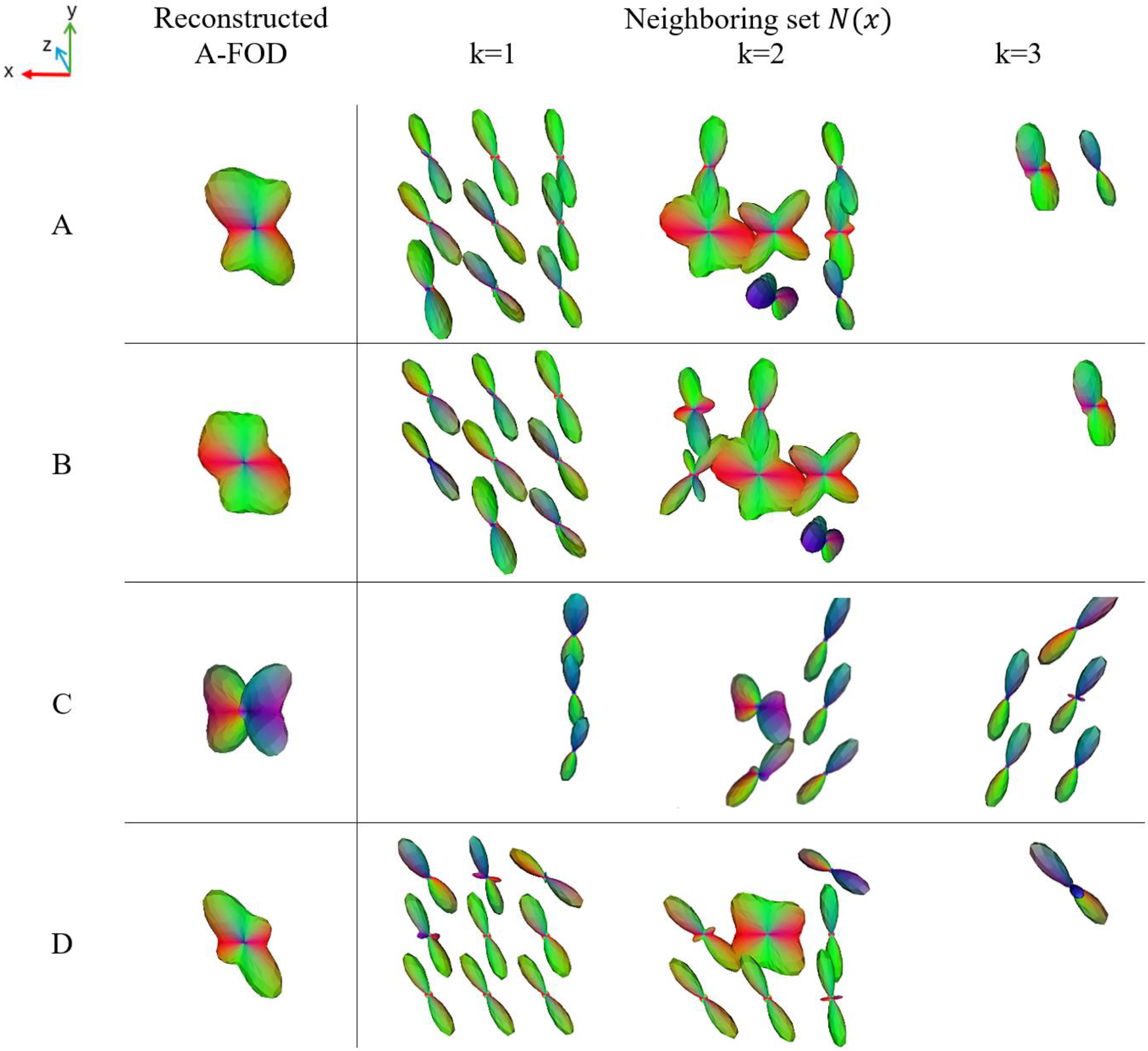
Characterization of EnCAR limitation in the DiSCo Phantom. Reliance on high-variance FODs and truncated neighboring sets *N*(*x*) (right) results in reconstructed A-FODs (left) that appear as blurred glyphs with poorly separable lobes. *N*(*x*) is visualized as a cubic 3 × 3 × 3 set decomposed by its depth index k (sub-columns k=1, 2, 3). The symmetric FODs were derived using MSMT-CSD.

In all samples illustrated in Fig. C.1, the neighboring sets are truncated due to boundary conditions. Furthermore, in samples A, B and D, the middle slices of the neighboring sets contain high-variance FODs. This uncertainty and lack of reliable information cause the model to suffer a significant loss of directional context. Without accurate and reliable FODs in *N*(*x*) to guide the self-supervised reconstruction, the resulting A-FODs appear blurred and lack clearly separable lobes.

## References

1. Basser, P.J., J. Mattiello, and D. LeBihan, Estimation of the effective self-diffusion tensor from the NMR spin echo. Journal of Magnetic Resonance, Series B, 1994. 103(3): p. 247–254.

2. Bastiani, M., et al., Improved tractography using asymmetric fibre orientation distributions. Neuroimage, 2017. 158: p. 205–218.

3. Bastiani, M. and A. Roebroeck, Unraveling the multiscale structural organization and connectivity of the human brain: the role of diffusion MRI. Frontiers in neuroanatomy, 2015. 9: p. 77.

4. Karayumak, S.C., E. Özarslan, and G. Unal, Asymmetric orientation distribution functions (AODFs) revealing intravoxel geometry in diffusion MRI. Magnetic Resonance Imaging, 2018. 49: p. 145–158.

5. Feng, Y. and J. He, Asymmetric fiber trajectory distribution estimation using streamline differential equation. Medical image analysis, 2020. 63: p. 101686.

6. Schilling, K.G., et al., Histologically derived fiber response functions for diffusion MRI vary across white matter fibers—An ex vivo validation study in the squirrel monkey brain. NMR in Biomedicine, 2019. 32(6): p. e4090.

7. Kjer, H.M., et al., Bridging the 3D geometrical organisation of white matter pathways across anatomical length scales and species. Elife, 2025. 13: p. RP94917.

8. Poirier, C. and M. Descoteaux, A unified filtering method for estimating asymmetric orientation distribution functions. NeuroImage, 2024. 287: p. 120516.

9. Wu, Y., et al., Mitigating gyral bias in cortical tractography via asymmetric fiber orientation distributions. Medical image analysis, 2020. 59: p. 101543.

10. Ehricke, H.-H., K.-M. Otto, and U. Klose, Regularization of bending and crossing white matter fibers in MRI Q-ball fields. Magnetic resonance imaging, 2011. 29(7): p. 916–926.

11. Bastiani, M., et al., Improved tractography by modelling sub-voxel fibre patterns using asymmetric fibre orientation distributions. Proc. Int. Soc. Magn. Reson. Med. Presented at the ISMRM. ISMRM, 2016.

12. Barmpoutis, A., et al. Extracting tractosemas from a displacement probability field for tractography in DW-MRI. in International Conference on Medical Image Computing and Computer-Assisted Intervention. 2008. Springer.

13. Meng, J., J. He, and Y. Wang, Estimation of fiber orientation distributions in superficial white matter using an asymmetric constrained spherical deconvolution method. Journal of Neuroscience Methods, 2025. 415: p. 110353.

14. Rafael-Patino, J., et al., The diffusion-simulated connectivity (DiSCo) dataset. Data in Brief, 2021. 38: p. 107429.

15. Tong, Q., et al., Multicenter dataset of multi-shell diffusion MRI in healthy traveling adults with identical settings. Scientific Data, 2020. 7(1): p. 157.

16. Liang, K.K., Efficient conversion from rotating matrix to rotation axis and angle by extending Rodrigues’ formula. arXiv preprint 1810.02999, 2018.

17. Vaswani, A., et al., Attention is all you need. Advances in neural information processing systems, 2017. 30.

18. Han, K., et al., A survey on vision transformer. IEEE transactions on pattern analysis and machine intelligence, 2022. 45(1): p. 87–110.

19. Descoteaux, M., High angular resolution diffusion MRI: from local estimation to segmentation and tractography. 2008, Université Nice Sophia Antipolis.

20. Tournier, J.-D., et al., Direct estimation of the fiber orientation density function from diffusion-weighted MRI data using spherical deconvolution. Neuroimage, 2004. 23(3): p. 1176–1185.

21. Raj, A., C. Hess, and P. Mukherjee, Spatial HARDI: improved visualization of complex white matter architecture with Bayesian spatial regularization. Neuroimage, 2011. 54(1): p. 396–409.

22. Devlin, J., et al. Bert: Pre-training of deep bidirectional transformers for language understanding. in Proceedings of the 2019 conference of the North American chapter of the association for computational linguistics: human language technologies, volume 1 (long and short papers). 2019.

23. Dosovitskiy, A., et al., An image is worth 16×16 words: Transformers for image recognition at scale. arXiv preprint 2010.11929, 2020.

24. Li, J., et al. Blip: Bootstrapping language-image pre-training for unified vision-language understanding and generation. in International conference on machine learning. 2022. PMLR.

25. Radford, A., et al. Learning transferable visual models from natural language supervision. In International conference on machine learning. 2021. PmLR.

26. Vivek, S., et al., Explainable variational autoencoder (E-VAE) model using genome-wide SNPs to predict dementia. Journal of biomedical informatics, 2023. 148: p. 104536.

27. Liu, W., et al. Towards visually explaining variational autoencoders. in Proceedings of the IEEE/CVF conference on computer vision and pattern recognition. 2020.

28. Gautam, S., et al., Protovae: A trustworthy self-explainable prototypical variational model. Advances in Neural Information Processing Systems, 2022. 35: p. 17940–17952.

29. Caruyer, E., et al., Design of multishell sampling schemes with uniform coverage in diffusion MRI. Magnetic resonance in medicine, 2013. 69(6): p. 1534–1540.

30. Kellner, E., et al., Gibbs-ringing artifact removal based on local subvoxel-shifts. Magnetic resonance in medicine, 2016. 76(5): p. 1574–1581.

31. Veraart, J., et al., Denoising of diffusion MRI using random matrix theory. Neuroimage, 2016. 142: p. 394–406.

32. Andersson, J.L., S. Skare, and J. Ashburner, How to correct susceptibility distortions in spinecho echo-planar images: application to diffusion tensor imaging. Neuroimage, 2003. 20(2): p. 870–888.

33. Andersson, J.L. and S.N. Sotiropoulos, An integrated approach to correction for off-resonance effects and subject movement in diffusion MR imaging. Neuroimage, 2016. 125: p. 1063–1078.

34. Zhai, F., et al., Disrupted white matter integrity and network connectivity are related to poor motor performance. Scientific Reports, 2020. 10(1): p. 18369.

35. Zhang, X., et al., Characterization of white matter changes along fibers by automated fiber quantification in the early stages of Alzheimer’s disease. NeuroImage: Clinical, 2019. 22: p. 101723.

36. Jeurissen, B., et al., Multi-tissue constrained spherical deconvolution for improved analysis of multi-shell diffusion MRI data. NeuroImage, 2014. 103: p. 411–426.

37. Tournier, J.-D., et al., MRtrix3: A fast, flexible and open software framework for medical image processing and visualisation. Neuroimage, 2019. 202: p. 116137.

38. Jeurissen, B., et al., Diffusion MRI fiber tractography of the brain. NMR in Biomedicine, 2019. 32(4): p. e3785.

39. Schilling, K.G., et al., Functional tractography of white matter by high angular resolution functional-correlation imaging (HARFI). Magnetic resonance in medicine, 2019. 81(3): p. 2011–2024.

40. Schilling, K.G., et al., Prevalence of white matter pathways coming into a single white matter voxel orientation: The bottleneck issue in tractography. Human brain mapping, 2022. 43(4): p. 1196–1213.

41. Tan, Y.-F., et al., Anatomy-to-tract mapping infers white matter pathways without diffusion streamline propagation. Nature Communications, 2025. 17: p. 36.

42. Khan, A., et al., A recent survey of vision transformers for medical image segmentation. IEEE Access, 2025.

43. Halder, A., et al., Implementing vision transformer for classifying 2D biomedical images. Scientific Reports, 2024. 14(1): p. 12567.

44. Zheng, W., et al., Lightweight transformer image feature extraction network. PeerJ Computer Science, 2024. 10: p. e1755.

45. Yi, S., et al., Spin-weighted spherical harmonics for polarized light transport. ACM Transactions on Graphics (TOG), 2024. 43(4): p. 1–24.

46. De Luca, A., et al., Cross-site harmonization of multi-shell diffusion MRI measures based on rotational invariant spherical harmonics (RISH). NeuroImage, 2022. 259: p. 119439.

47. Yan, H., et al., Estimating fiber orientation distribution from diffusion MRI with spherical needlets. Medical image analysis, 2018. 46: p. 57–72.

48. Zhang, D., et al., Asymmetric Fiber Orientation Distribution Estimation and Response Function Calibration with an Unsupervised Deconvolution Network.

49. Hendriks, T., A. Vilanova, and M. Chamberland, Implicit neural representation of multi-shell constrained spherical deconvolution for continuous modeling of diffusion MRI. Imaging Neuroscience, 2025. 3: p. imag_a_00501.

50. Rheault, F., et al., Bundle-specific tractography with incorporated anatomical and orientational priors. Neuroimage, 2019. 186: p. 382–398.

51. Consagra, W., L. Ning, and Y. Rathi, Neural orientation distribution fields for estimation and uncertainty quantification in diffusion MRI. Medical Image Analysis, 2024. 93: p. 103105.

52. Maaten, L.v.d. and G. Hinton, Visualizing data using t-SNE. Journal of machine learning research, 2008. 9(Nov): p. 2579–2605.

